# N-Acetylglucosamine ameliorates knee osteoarthritis by enhancing UDP-GlcNAc conversion and chondrocyte DNA repair capacity

**DOI:** 10.64898/2026.10.21.760584

**Authors:** Changze Song, Meilu Li, Xiangfei Qi, Huisheng Yuan, Haohan Zhang, Jinghan Zhang, Hongyin Huang, Zhenghao Li, Junhao Chu, Xiaohan Wang, Yong Yang, Tianzhi Zhang, Chunxia Zhang, Zilong Wang

## Abstract

**Background:** Knee osteoarthritis (KOA) is a degenerative joint disease characterized by progressive cartilage degradation and synovial inflammation, while current therapies offer only symptomatic relief. This study aimed to determine whether N-acetylglucosamine (GlcNAc) is superior to glucosamine hydrochloride (GlcN·HCl) in slowing the progression of KOA and to elucidate the underlying mechanisms involved.

**Methods:** Liquid chromatography-tandem mass spectrometry (LC-MS/MS) was employed to assess serum concentrations of UDP-GlcNAc to conduct a comprehensive evaluation of the pharmacokinetics and pharmacodynamics associated with both GlcNAc and GlcN·HCl. Using a destabilization of the medial meniscus (DMM) rat model, we performed micro-computed tomography and histopathological assessments to evaluate structural changes in the joint tissues. Additionally, we conducted bulk RNA sequencing, flow cytometry and ELISA to validate the efficacy of GlcNAc in anti-inflammatory immunoregulation and DNA repair in knee cartilage.

**Results:** GlcNAc achieved higher peak serum concentrations and more efficient conversion to UDP-GlcNAc than GlcN·HCl, with UDP-GlcNAc levels 4.57 times higher at 9 hours after administration. GlcNAc significantly attenuated KOA-like pathological changes, downregulated catabolic markers *Mmp3* and *Adamts4*, decreased pain-related neural markers *Pgp9*.*5* and *Ngf*, and alleviated pain sensitivity. RNA-seq identified 4,395 downregulated DEGs in DMM rats, of which GlcNAc upregulated 2,808 compared to only 391 by GlcN·HCl. GlcNAc promoted DNA repair and adaptive immune cell differentiation pathways, and decreased serum IL-1β, IL-6, and IL-17α. Four candidate DNA repair genes, *Hist1h2an, Poli, Ube2t*, and *Hells*, were identified as significantly upregulated by GlcNAc, with GSEA confirming homologous recombination repair pathway enrichment.

**Conclusion:** This study shows GlcNAc outperforms GlcN·HCl in rat DMM knee osteoarthritis via multiple mechanisms, including 4.57-fold higher UDP-GlcNAc conversion, reduced cartilage degeneration, immune rebalancing, and DNA repair activation, supporting GlcNAc as a promising KOA candidate warranting further clinical study.

## Introduction

Knee osteoarthritis (KOA) is a degenerative knee joint disease characterized by gradual articular cartilage degradation and subchondral bone damage, resulting in joint stiffness, chronic joint pain, and even physical disability for a significant portion of the population. KOA affects approximately 7% of the world’s population and has an estimated prevalence of 73% worldwide in people older than 55 years of age. In recent years, the prevalence of KOA has increased among populations as young as 30, imposing substantial clinical, social, and economic burdens globally^[1]^. Conservative therapy is recommended for symptomatic early-stage osteoarthritis of the knee, and surgery, including knee replacement, for late-stage disease or failure of conservative therapies^[2]^. Currently, multiple conservative therapies for KOA are employed in clinical practice, such as weight control, exercise, physical therapy (including acupuncture, laser therapy, and electromagnetic therapy), oral analgesics, and intra-articular injections^[3]^. However, given that current conservative treatment options have limited long-term efficacy and are associated with an increased risk of gastrointestinal, cardiovascular, and renal adverse effects^[4]^, the management of KOA remains challenging. Therefore, investigating a critical agent to inhibit cartilage degeneration holds important clinical implications.

Glucosamine (GlcN), as a natural amino acid monosaccharide, is considered the main building block for the glycosaminoglycan structure of cartilage found in the joints^[5]^. Glucosamine sulfate (GlcN·H2SO4) and glucosamine hydrochloride (GlcN·HCl) are the main nutraceuticals for cartilage. However, the published results of the Glucosamine/chondroitin Arthritis Intervention Trial (GAIT) showed that, in the overall group of patients with KOA, GlcN·HCl alone was not significantly more effective than placebo in reducing pain^[6]^. Similarly, in mouse models of KOA, an excessive amount of GlcN·H2SO4 administration significantly accelerated the progression of KOA, as evidenced by augmented cartilage destruction and accelerated KOA-associated bone remodeling in the subchondral trabecular bone^[7]^. Thus, there is an urgent need for alternative glucosamine-based strategies to facilitate more effective treatment of KOA.

N-acetylglucosamine (GlcNAc) is an acetylated derivative of glucosamine and is the fundamental substrate of many polysaccharides within biological cells. The molecular weight of GlcNAc is only 221.21 Da^[8]^, permitting it to pass through cell membranes to exert biological functions, such as antioxidant, and anti-inflammatory effects^[9]^. However, whether GlcNAc can confer a protective effect on knee cartilage is currently unclear.

In this study, we aimed to investigate the protective effect of GlcNAc on KOA through in vivo KOA mouse models. It lays a theoretical foundation for the development of GlcNAc-based dietary supplements for the alleviation of KOA.

## Materials and methods

### Reagents

N-acetylglucosamine (Rolvanferm®, Purity: 99.35%, Lot No. RH202502019) was produced by Shandong Runhan Biotechnology Co., Ltd. and prepared by fermentation of Bacillus subtilis BNZR2-7. Glucosamine hydrochloride (GPC standards, Purity: 99.97%, Lot No. LS505784) was purchased from Shanghai Yuanye Biotechnology Co., Ltd.

### Knee osteoarthritis rats model establishment

We assessed the therapeutic efficacy of GlcNAc or GlcN·HCl in a rat model of KOA. The KOA model was induced by destabilization of the medial meniscus (DMM) surgery, as described previously. Briefly, animals were anesthetized with 2%–3% isoflurane. The bilateral knee areas were disinfected with 0.5% iodophor, after which the hindlimb was positioned at 90° flexion and immobilized. A 3-mm longitudinal incision was made along the medial border of the patellar ligament. Blunt dissection was then performed to separate the subcutaneous tissues. The joint capsule was then opened medially to expose the medial meniscotibial ligament (MMTL), which was subsequently severed with micro-scissors. Successful joint destabilization was verified during the procedure. The surgical site was flushed with sterile saline, subsequently closing the joint capsule and skin. In the sham-operated group, the same surgical approach was used, except that the MMTL remained undisturbed.

### GlcNAc or GlcN·HCl oral administration

For the surgical study, a total of 32 rats were randomly assigned to four experimental groups: a sham-operated control group (Sham), a group subjected to destabilization of the medial meniscus (DMM), a DMM group treated with GlcN·HCl (DMM + GlcN·HCl), and a DMM group receiving GlcNAc (DMM + GlcNAc), with 8 animals per group. During the 60-day study period, all groups performed daily climbing exercise in the same animal housing environment as previously described. At the end of the experiment, three seven-week-old SPF-grade Sprague-Dawley rats were additionally selected and left untreated as the untreated control group (WT).

### Micro-CT analysis of live rats

Inhaled diethyl ether was used for brief anesthesia during micro-CT scanning. Micro-CT imaging was used to scan the samples, with the system configured to operate at a voltage of 90 kV and a current of 70 μA, yielding a spatial resolution of 10 μm. A uniform threshold was applied across all groups during image analysis, ensuring consistent three-dimensional reconstruction of the specimens. For histomorphometric evaluation, the region extending from the proximal tibial cartilage to the area beneath the growth plate was delineated and subjected to three-dimensional analysis.

### Histological analysis

For histological assessment, knee joint specimens were initially fixed in a 4% paraformaldehyde solution for 72 hours, followed by decalcification in EDTA for three weeks and then paraffin embedding. Coronal sections, each 4 μm thick, were subjected to hematoxylin-eosin and Alcian Blue staining to facilitate morphological evaluation. The extent of osteoarthritis-like pathology was quantified using multiple grading systems, including the OARSI scale, Mankin criteria, synovitis index, and measurements of cartilage area. To ensure reliability, two independent observers assessed the severity of the KOA-like phenotype utilizing the OARSI scoring method on three distinct sections per joint, specifically focusing on the medial femoral condyle and medial tibial plateau.

### Bulk RNA-seq and data analyses of knee cartilage

Cartilage RNA extraction was carried out using the highly efficient Trizol-based protocol. In brief, the cartilage tissue was gently disrupted with a tissue grinding rod for 16-25 strokes, followed by lysis in 500 μL of Trizol reagent (Invitrogen). After incubation at room temperature and subsequent freezing in liquid nitrogen, the samples were preserved at -80 °C. RNA isolation was performed through chloroform-mediated phase separation, followed by precipitation with isopropanol and two successive washes with 75% ethanol. The resulting RNA pellet was air-dried and reconstituted in 20 μL of RNase-free ddH2O for subsequent applications. Sequencing libraries were constructed using the Illumina TruSeq Stranded mRNA LT Sample Prep Kit in accordance with the manufacturer’s instructions. Briefly, mRNA was enriched from total RNA using oligo(dT)-coated magnetic beads. The enriched RNA was then fragmented and reverse-transcribed into cDNA. The resulting cDNA fragments were subjected to end repair, adenylation, and ligation with Illumina adapters. Following PCR enrichment, libraries were quantified and quality-assessed prior to sequencing. Paired-end sequencing was performed on the Illumina HiSeq platform. Differential expression analysis was conducted using the DESeq2 package in R (version 4.2.1). Genes meeting the thresholds of adjusted p-value < 0.05 and |log2 fold change| > 1 were considered statistically significant.

### Flow cytometry assay

Single-cell suspensions were prepared from the bone marrow of KOA rats. To evaluate surface marker expression, cells were incubated with antibodies in PBS supplemented with 1% (w/v) BSA for 30 minutes on ice. The staining panel consisted of Super Bright 600-conjugated anti-CD4 (Clone: SK-3; eBioscience), BV510-conjugated anti-CD8a (Clone: 53-6.7; BioLegend), APC-Cy7-conjugated anti-TCRβ (Clone: H57-597; BioLegend), and BV605-conjugated anti-B220 (Clone: RA3-6B2; BioLegend). Flow cytometric acquisition was conducted using the Beckman CytoFLEX S. Data analysis was performed using FlowJo software.

### Enzyme-linked immunosorbent assay

Using ELISA kits from Lianke Biotech, the concentrations of IL-1β, IL-6, and IL-17α in rat serum were detected. Cell culture supernatants from each group were collected. The ELISA plate was soaked, washing solution was added and left to stand for 30 seconds, then the washing solution was discarded. 50 μL of standard substance, culture medium, or sample was added to each well, the plate was sealed with sealing film, and incubated at 37°C for 2 hours. The liquid was discarded, 300 μL of washing solution was added to each well, and this washing step was repeated 5-6 times. 100 μL of horseradish peroxidase was added to each well, the plate was sealed with new sealing film, and incubated at 37°C for 45 minutes. Washing solution was applied, and this process was repeated 5-6 times. 100 μL of chromogenic substrate TMB was added to each well, and incubated at 37°C in the dark for 20-30 minutes. 100 μL of stop solution was added to each well, and the reaction color changed from blue to yellow.

### Liquid chromatography-tandem mass spectrometry (LC-MS/MS) system

Under aseptic conditions, draw whole blood and inject it into a dry test tube/centrifuge tube without anticoagulant to prevent hemolysis. Place the tube vertically at room temperature for 30-60 minutes (or accelerate the process at 37°C) until the blood fully coagulates and the clot contracts. Centrifuge at 3000 rpm for 15-30 minutes to sediment the blood cells and clots. Carefully aspirate the upper clear liquid with a sterile pipette, avoiding contact with the precipitate, and then sub - aliquot it into sterile tubes.

Quantification of GlcNAc, GlcN·HCl and UDP-GlcNAc were performed using a LC-MS/MS system consisting of an AB SCIEX 6500 mass spectrometer equipped with an electrospray ionization (ESI) source operated in negative ion mode. Chromatographic Conditions: Chromatographic separation was achieved on a Phenomenex Luna® HILIC column (5 μm, 150 × 3 mm) maintained at 45°C. The mobile phase consisted of (A) 2 mM ammonium acetate in water and (B) acetonitrile. The flow rate was set at 0.8 mL/min with the following gradient program: 0– 2 min, 50% B. The autosampler temperature was maintained at 6°C, and the injection volume was 10.0 μL. Parameters: Ion Spray Voltage -4500 V; Temperature 550°C; GS1 40 psi; GS2 40 psi; CUR 35 psi; CAD 10 psi. Declustering Potential (DP) -60 V, -30 V (IS); Collision Energy (CE) - 35 V, –25 V (IS); Cell Exit Potential (CXP) -10 V; Entrance Potential (EP) -10 V. The dwell time for each channel was set at 100 ms.

### Statistical analysis

The statistical analyses were performed using the R programming language (v4.2.1) and the figures were generated through the ggplot2 package. The pod diagram was analyzed through the stats and car packages, and the Venn diagrams were generated by the VennDiagram package. The comparative analyses and correlation analyses were performed by t-test and Spearman’s correlation test, respectively. K-Viewer (1.7.1.1) and OlyVIA (V4.1) were used for histological scanning image system. In this study, bulk RNA-Seq differential genes |Log2FC|> 1 and P adj. < 0.05 were statistically significant.

## Results

### The pharmacokinetics and pharmacodynamics of GlcNAc and GlcN·HCl

To investigate the pharmacokinetics and pharmacodynamics of GlcNAc and GlcN·HCl, we administered GlcNAc or GlcN·HCl orally to normal rats and measured serum levels of UDP-GlcNAc and the respective compound at designated time points (**Figure 1A**). For GlcNAc, the peak concentration reached 3071.67 ng/mL, and the time to peak occurred at almost 2 hours, with a half-life of approximately 3 hours (**Figure 1B**). The concentration returned to baseline (286.00 ng/mL) after more than 6 hours have elapsed, and the AUC0-6 was calculated to be 7505.73 ng·h/mL (**Figure 1B**).

**Figure 1.**
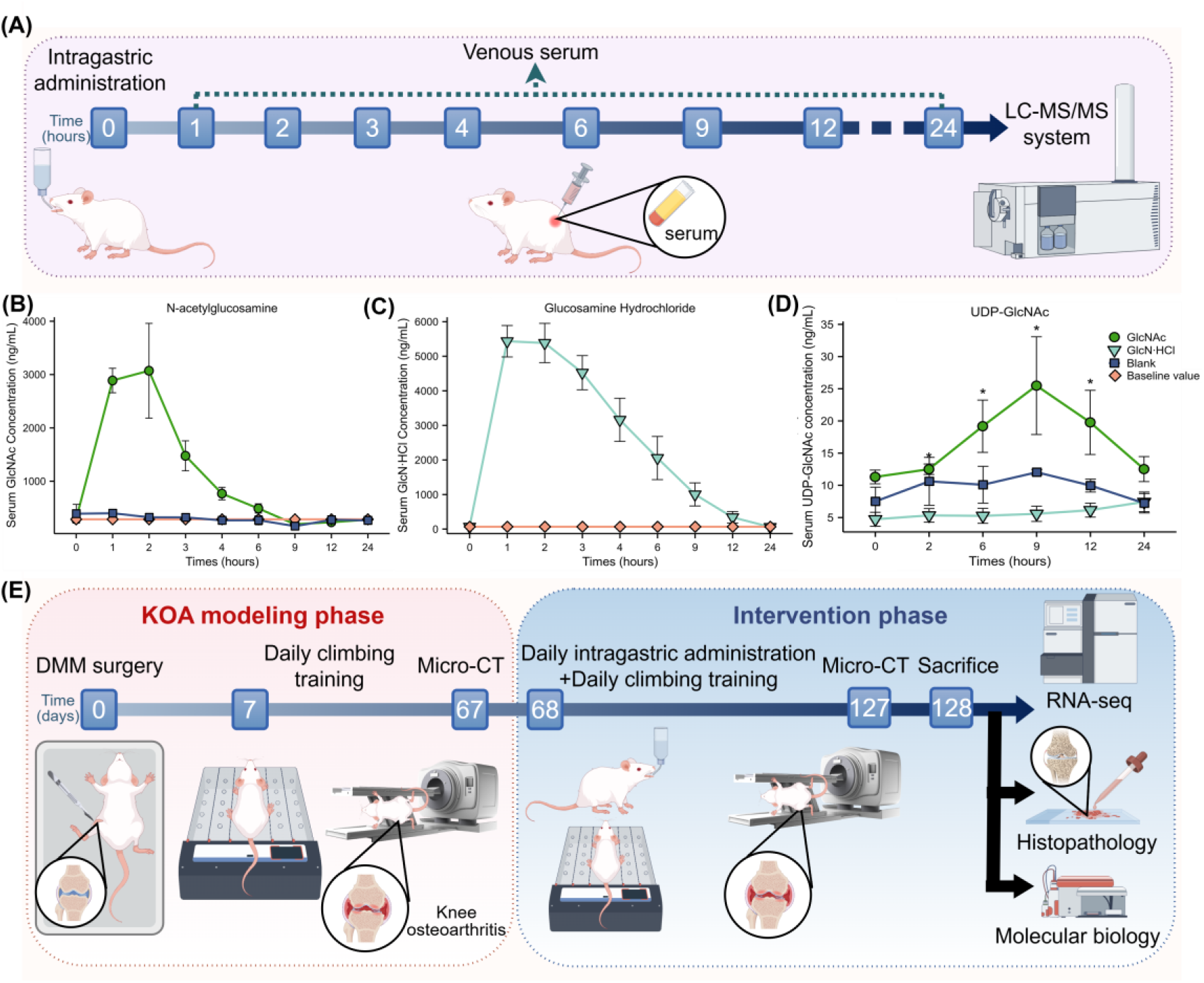
The schematic diagram of this study and the pharmacokinetics and pharmacodynamics of GlcNAc and GlcN·HCl. (A) The schematic diagram of the pharmacokinetics and pharmacodynamics study. (B,C) The pharmacokinetics of GlcNAc (B) and GlcN·HCl (C). (D) The pharmacodynamics of GlcNAc and GlcN·HCl based on UDP-GlcNAc. (E) The schematic diagram of the pharmacological intervention study.

For GlcN·HCl, the peak concentration reached 5435.00 ng/mL, and the time to peak was approximately 1 hour, with a half-life of approximately 4 hours (**Figure 1C**). The concentration returned to baseline (67.78 ng/mL) after more than 12 hours had elapsed, and the AUC0-12 was calculated as 27960.41 ng·h/mL (**Figure 1C**).

Both N-acetylglucosamine and Glucosamine Hydrochloride must ultimately be converted into uridine diphosphate-N-acetylglucosamine (UDP-GlcNAc) in vivo to exert their corresponding biological effects. Therefore, in this study, UDP-GlcNAc was used as the common metabolite of GlcNAc and GlcN·HCl to investigate the pharmacodynamic properties of the two substances. Orally administered GlcNAc begins converting into UDP-GlcNAc within 2 hours. The concentration of UDP-GlcNAc peaks at 9 hours, with an average concentration of 25.48 ng/mL, which is 4.57 times that of GlcN·HCl (**Figure 1D**). At 24 hours after oral administration, UDP-GlcNAc is completely metabolized. Glucosamine hydrochloride exhibits a notably low conversion rate. Over the 24 hours following oral administration, no significant variation was observed, and the mean conversion concentration at 9 hours post-administration amounted to just 5.57 ng/mL (**Figure 1D**).

### GlcNAc decelerates KOA progression in rats with DMM

To compare the effect of GlcNAc with GlcN·HCl on the progression of KOA in rats, the DMM surgery was performed to generate the KOA rats, designated as the DMM group. GlcNAc or GlcN·HCl was then intragastrically administered after 7-day recovery periods and 60 days of daily climbing training following DMM surgery. Micro-computed tomography (CT) imaging, histopathological evaluation, and cartilage metabolic biomarkers were employed to assess the progression of KOA and the efficacy of drug treatments (**Figure 1E**).

Then, we performed micro-CT, histomorphometric, and histopathological analyses and bulk RNA sequencing to evaluate the differential therapeutic effects of GlcNAc and GlcN·HCl at the end of the 60-day treatment period (127 days post-DMM surgery). DMM + GlcNAc rats exhibited a marked attenuation of KOA-like pathological changes, compared with DMM and DMM + GlcN·HCl groups (**Figure 2A**). For instance, we observed a significant reduction in the Kellgren-Lawrence grade on micro-CT imaging (**Figure 2A**), the ICRS grade from histomorphometric analyses (**Figure 2B**), and the OARSI score, Mankin’s score, and synovitis score (**Figure 2C**) from histopathological analyses, alongside an increase in cartilage area.

**Figure 2.**
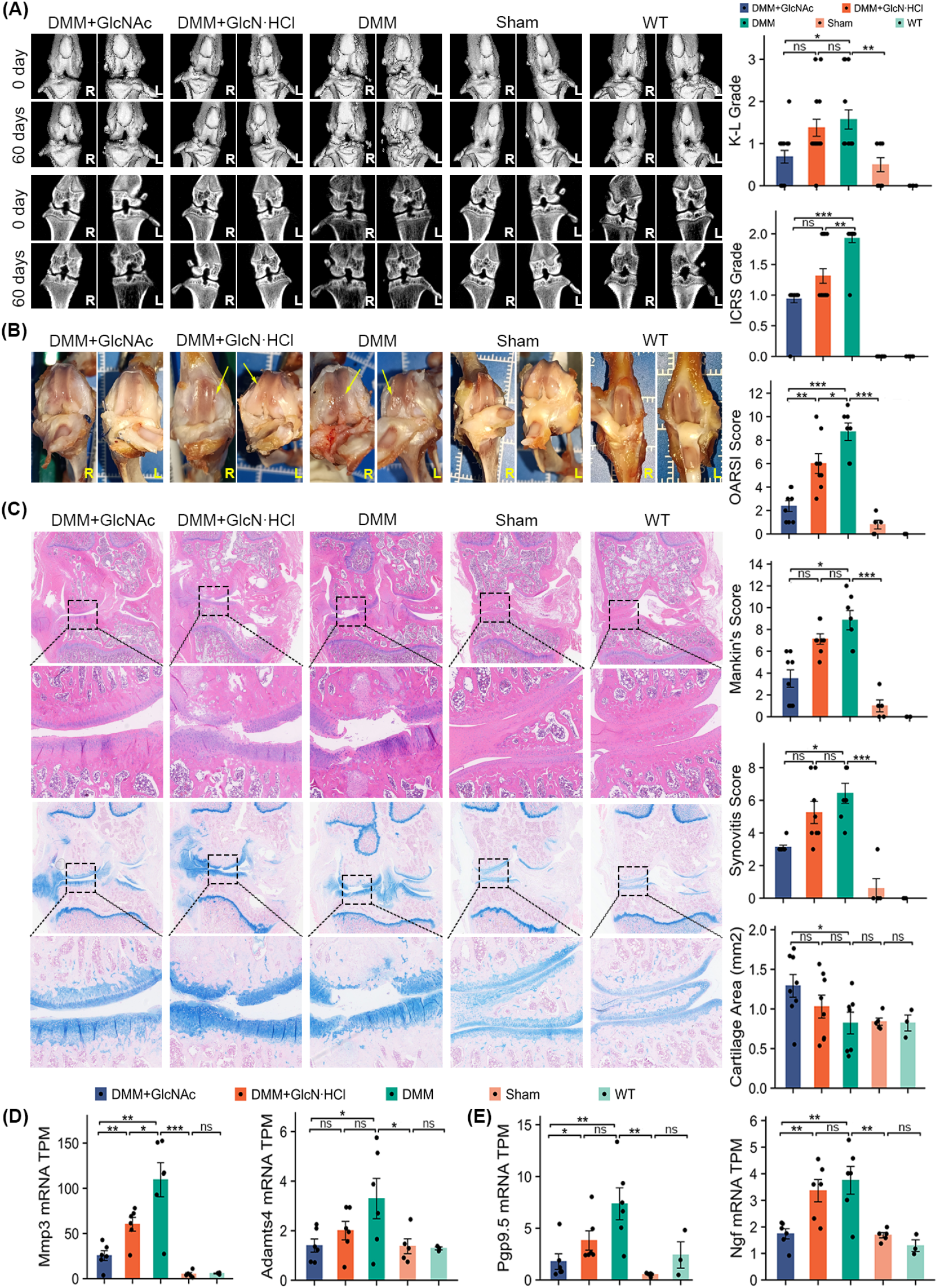
GlcNAc ameliorates cartilage degradation and the pain in KOA rats. (A) Representative micro-CT images of live sham-or DMM-operated WT rats provided with GlcNAc or GlcN·HCl or ddH2O. (B) The gross pathologic examination of knee cartilage of live sham-or DMM-operated WT rats provided with GlcNAc or GlcN·HCl or ddH2O. (C) The histological and histomorphometric analysis of cartilage showed chondroprotective effects of GlcNAc or GlcN·HCl. (D) Bulk RNA sequencing analyses and quantification of catabolic marker genes (*Mmp3* and *Adamts4*) indicated the protective effect of GlcNAc on cartilage matrix under KOA. (E) Bulk RNA sequencing analyses and quantification of classical neural markers like *protein gene product 9*.*5* (*Pgp9*.*5*), *nerve growth factor* (*Ngf*), and the results confirmed the pain inhibition effect of GlcNAc.

Because GlcNAc showed excellent efficacy against KOA in imaging examinations and histological assessments, we aimed to examine the expression of cartilage metabolic markers and pain-related neural markers in the cartilage transcriptome data from the bulk RNA sequencing database. MMP3 and ADAMTS4 are the representative catabolic markers of knee cartilage cells and the results revealed a significant increase in the expression of the *Mmp3* and *Adamts4* genes in DMM rats, while GlcNAc treatment markedly reduced their expression compared with those in the DMM and DMM + GlcN·HCl groups (**Figure 2D**).

### GlcNAc relieves KOA pain

Pain is the primary reason OA patients seek medical care, making its alleviation crucial for assessing treatment efficacy. Significant decreases in pain sensitivity were observed in GlcNAc + DMM rats at the end of the 60-day treatment period, indicating that GlcNAc may effectively alleviate KOA-associated pain. To confirm the pain-reliving effects of GlcNAc at the molecular level, we examined expression of pain-related neurochemical marker genes in knee cartilage. Classical neural markers, such as *protein gene product 9*.*5* (*Pgp9*.*5*) and *nerve growth factor* (*Ngf*) were upregulated in DMM rats but were significantly downregulated after GlcNAc treatment (**Figure 2E**). These findings underscore GlcNAc’s potential to alleviate KOA-associated pain and improve activity levels in a rat model of KOA, which offers promising therapeutic implications.

### GlcNAc reduced immunoinflammatory response of cartilage in rats with KOA

To further explore the precise mechanism of GlcNAc on cartilage in KOA, we analyzed RNA-seq data from our laboratory rats (**Figure 3A**). Our RNA-seq identified 4,395 downregulated DEGs in DMM rats. Among these downregulated DMM-associated DEGs, GlcNAc upregulated 2,808 DEGs, while GlcN·HCl could only reverse the expression of 391 DEGs, 384 of which overlapped with those affected by GlcNAc (**Figure 3B**). Differential gene expression analysis identified significant upregulation of DNA repair and adaptive immune cell differentiation pathways in KOA rats treated with GlcNAc (**Figure 3C**).

**Figure 3.**
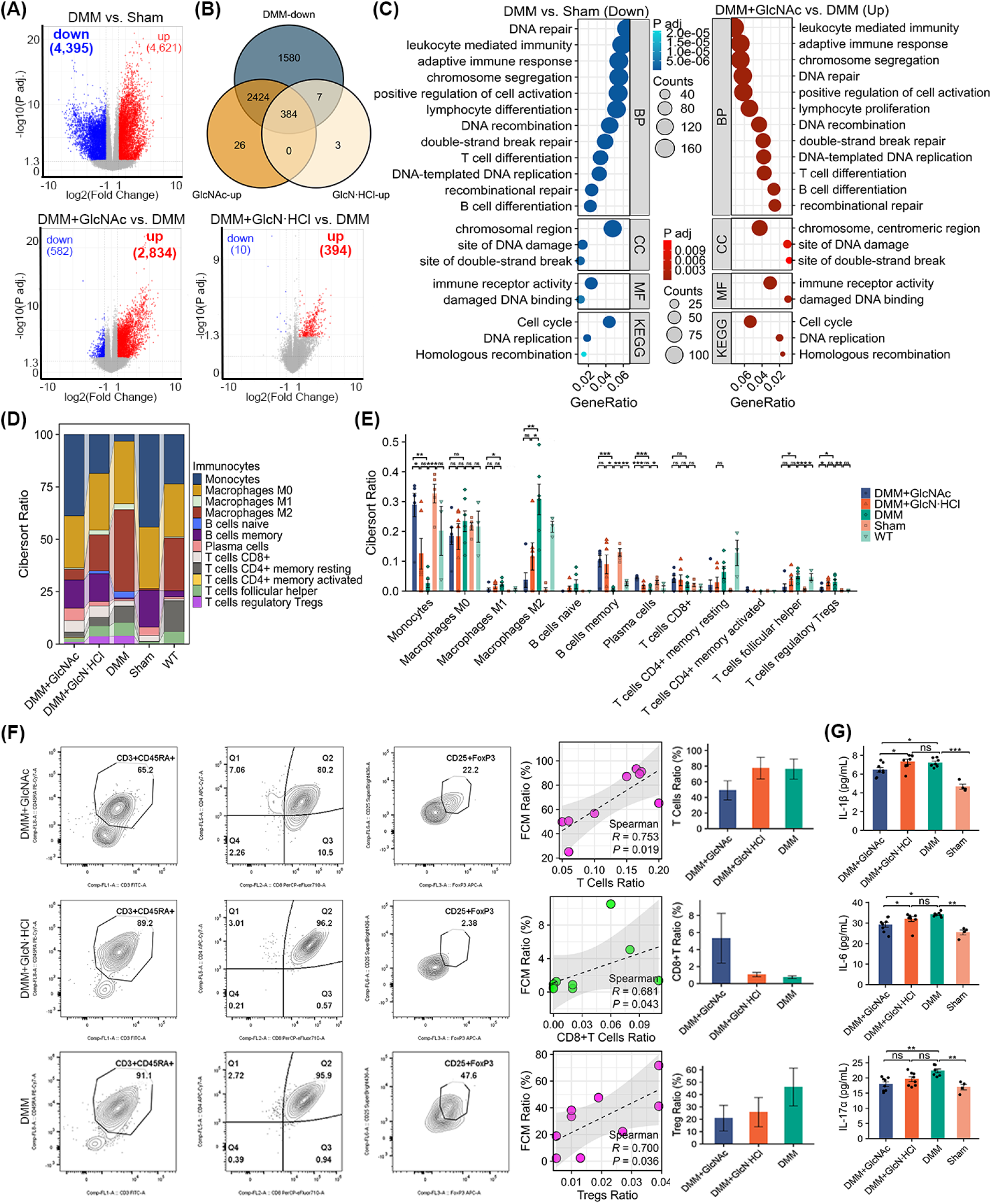
GlcNAc exerts its anti-inflammation immunoregulation in KOA rats. (A) The Volcano plot analysis of the differentially expressed genes (DEGs) in RNA-seq dataset. (B) Venn analysis of the downregulated DEGs in DMM set, the upregulated DEGs in GlcNAc dataset, and the upregulated DEGs in GlcN·HCl dataset. (C) Gene Ontology (GO) analysis of the downregulated DEGs in the DMM dataset and the upregulated DEGs in the GlcNAc dataset. (D,E) The immune infiltration analysis of RNA-seq dataset. (F) The flow cytometry about bone marrow of rats for immune infiltration analysis. (G) ELISA detected inflammatory factors IL-1β, IL-6 and IL-17α levels.

The immune infiltration analysis showed that GlcNAc had a low abundance of immunoinflammatory cells such as M1 macrophages, Tregs, and CD4+ T cells, whereas elevated levels of monocytes, plasma cells, CD8+ T cells and other non-inflammatory cells were observed (**Figure 3D,E**). To further validate the anti-inflammatory properties of GlcNAc in KOA rats, we performed flow cytometry (FCM) about bone marrow of rats. The findings were consistent with the bulk RNA sequencing results, which showed that Tregs exhibited downward trends and CD8+ T cells were elevated following GlcNAc administration (**Figure 3F**). The positive association between FCM ratio and immune cell ratio verified the consistency with the results of immune infiltration analysis based on bulk RNA sequencing (**Figure 3F**).

In order to explore the role of GlcNAc in alleviating inflammation more thoroughly, we subsequently measured serum levels of inflammatory factors using enzyme-linked immunosorbent assay (ELISA) in KOA rats. Consistent with the trend observed in the results from bulk RNA sequencing and FCM, ELISA results showed that GlcNAc reduced IL-1β, IL-6 and IL-17α levels, validating the anti-inflammatory properties of GlcNAc in KOA rats (**Figure 3G**).

### GlcNAc improves cartilage repair in rats with KOA

Whereas differential expression analysis of GO|KEGG and gene set enrichment analysis (GSEA) revealed significant upregulation of DNA repair in DMM + GlcNAc rats (**Figure 3C, 4A**), we found 91 DNA repair genes were upregulated in DMM + GlcNAc rats (**Figure 4B**). Using the Random Forests machine learning algorithm (**Figure 4C**), we pinpointed four candidate genes: *Hist1h2an, Poli, Ube2t*, and *Hells* (**Figure 4D**). Subsequent analysis revealed a statistically significant increase in the expression of these genes in DMM + GlcNAc rats compared with their expression in the DMM and DMM + GlcN·HCl groups (**Figure 4E**). Additionally, the GSEA enrichment analysis of the homologous recombination repair pathway corroborated the functions of these four genes (**Figure 4F**).

**Figure 4.**
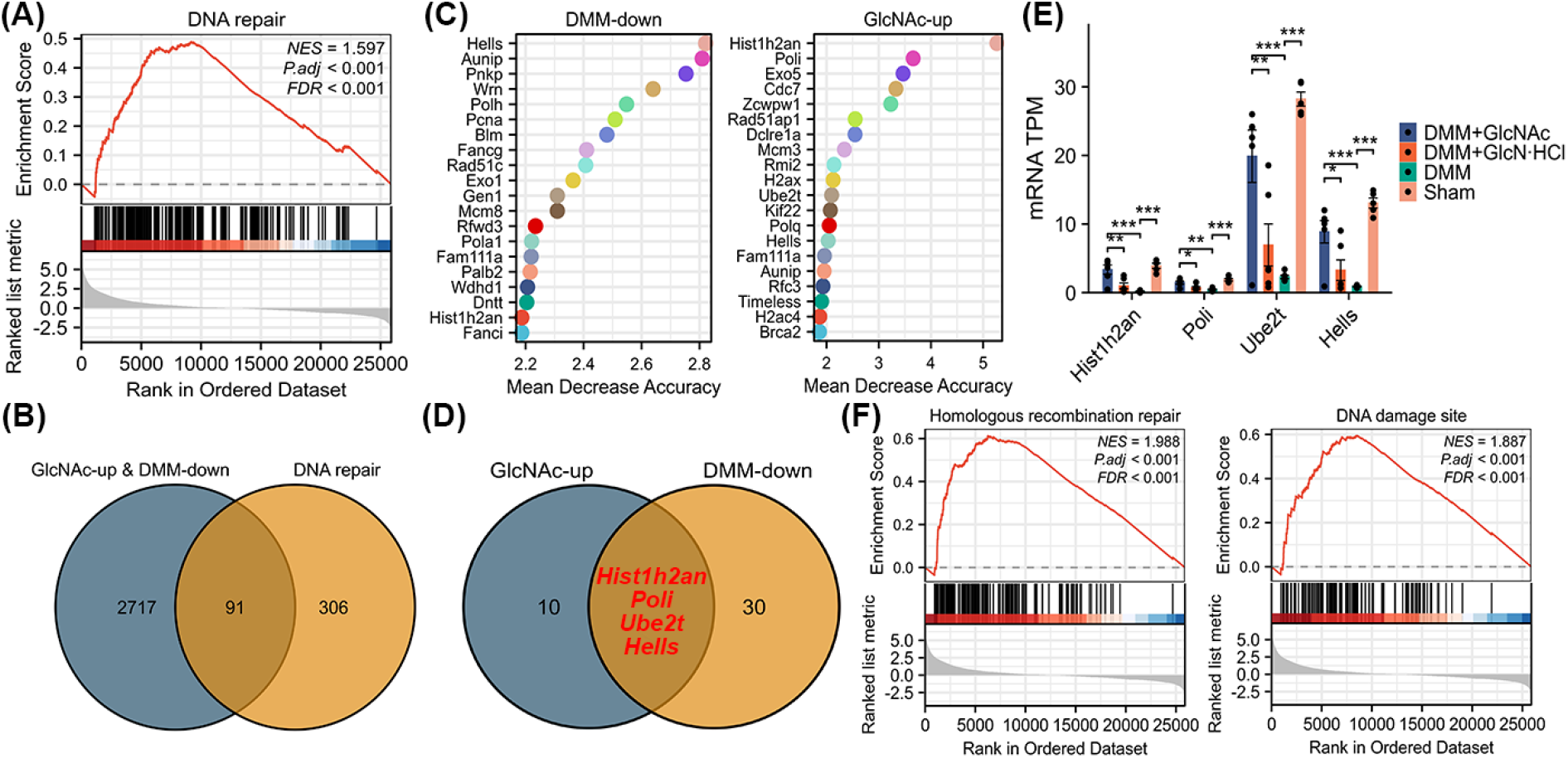
GlcNAc enhances DNA repair in knee cartilage of KOA rats through machine learning algorithm. (A) GSEA analysis of the upregulated DEGs of the GlcNAc dataset in the DNA repair pathway. (B) Venn analysis of the DNA-repair-related gene set, and the overlap DEGs between GlcNAc and DMM. (C) The Random Forests machine learning algorithm of the downregulated DEGs of the DMM dataset and the upregulated DEGs of the GlcNAc dataset. (D) Venn analysis of DEGs following Random Forests machine learning algorithm. (E) Comparison of identified four differential genes among live sham-or DMM-operated WT rats provided with GlcNAc or GlcN·HCl or ddH2O. (G) GSEA analysis of the upregulated DEGs of the GlcNAc dataset in the homologous recombination repair pathway validates the roles of these four identical differential genes.

## Discussion

Knee osteoarthritis (KOA) is a prevalent degenerative joint disorder characterized by progressive articular cartilage degradation, synovial inflammation, and subchondral bone remodeling, affecting a substantial proportion of the aging population worldwide and represents a leading cause of chronic pain and disability^[10, 11]^. Current therapeutic options, such as analgesics, nonsteroidal anti-inflammatory drugs, and glucosamine-based supplements, provide only symptomatic relief without halting disease progression^[4]^. Although GlcN·HCl is widely utilized as a dietary supplement for joint health, its clinical efficacy remains controversial^[6]^, owing to inefficient conversion to the active metabolite UDP-GlcNAc. In contrast, GlcNAc acts as a direct precursor in the hexosamine biosynthetic pathway and may represent a more efficient substrate for UDP-GlcNAc production. However, systematic comparative studies that evaluate the therapeutic superiority and underlying molecular and immunological mechanisms of GlcNAc versus GlcN·HCl in KOA remain lacking.

This study was therefore designed to address this critical gap by employing the DMM rat model of KOA and oral administration of GlcNAc or GlcN·HCl. A comprehensive multi-modal assessment was then conducted, integrating micro-CT imaging, histological grading, bulk RNA sequencing, immune infiltration analysis, flow cytometry, ELISA, and LC-MS/MS pharmacokinetic profiling. The specific objectives were to compare the pharmacokinetic profiles of GlcNAc and GlcN·HCl, evaluate their differential efficacy in decelerating KOA progression, and elucidate the mechanisms by which GlcNAc alleviates pain, reduces immunoinflammatory responses, and promotes cartilage repair through DNA repair pathway activation. Key findings demonstrated that GlcNAc achieved higher peak serum concentrations and more efficient conversion to UDP-GlcNAc than GlcN·HCl, with UDP-GlcNAc levels 4.57 times higher at 9 hours after administration. GlcNAc significantly attenuated KOA-like pathological changes, downregulated catabolic markers Mmp3 and Adamts4, decreased pain-related neural markers *Pgp9*.*5* and *Ngf*, and alleviated pain sensitivity. RNA-seq identified 4,395 downregulated DEGs in DMM rats, of which GlcNAc upregulated 2,808 compared to only 391 by GlcN·HCl. GlcNAc promoted DNA repair and adaptive immune cell differentiation pathways, reduced M1 macrophages, Tregs, and CD4+ T cells, elevated monocytes, plasma cells, and CD8+ T cells, and decreased serum IL-1β, IL-6, and IL-17α. Four candidate DNA repair genes, *Hist1h2an, Poli, Ube2t*, and *Hells*, were identified as significantly upregulated by GlcNAc, with GSEA confirming homologous recombination repair pathway enrichment (**Figure 5**). These findings collectively establish GlcNAc as a superior therapeutic candidate for KOA, with mechanisms that include enhanced UDP-GlcNAc concentration, anti-inflammatory immunomodulation, and promotion of cartilage DNA repair.

**Figure 5.**
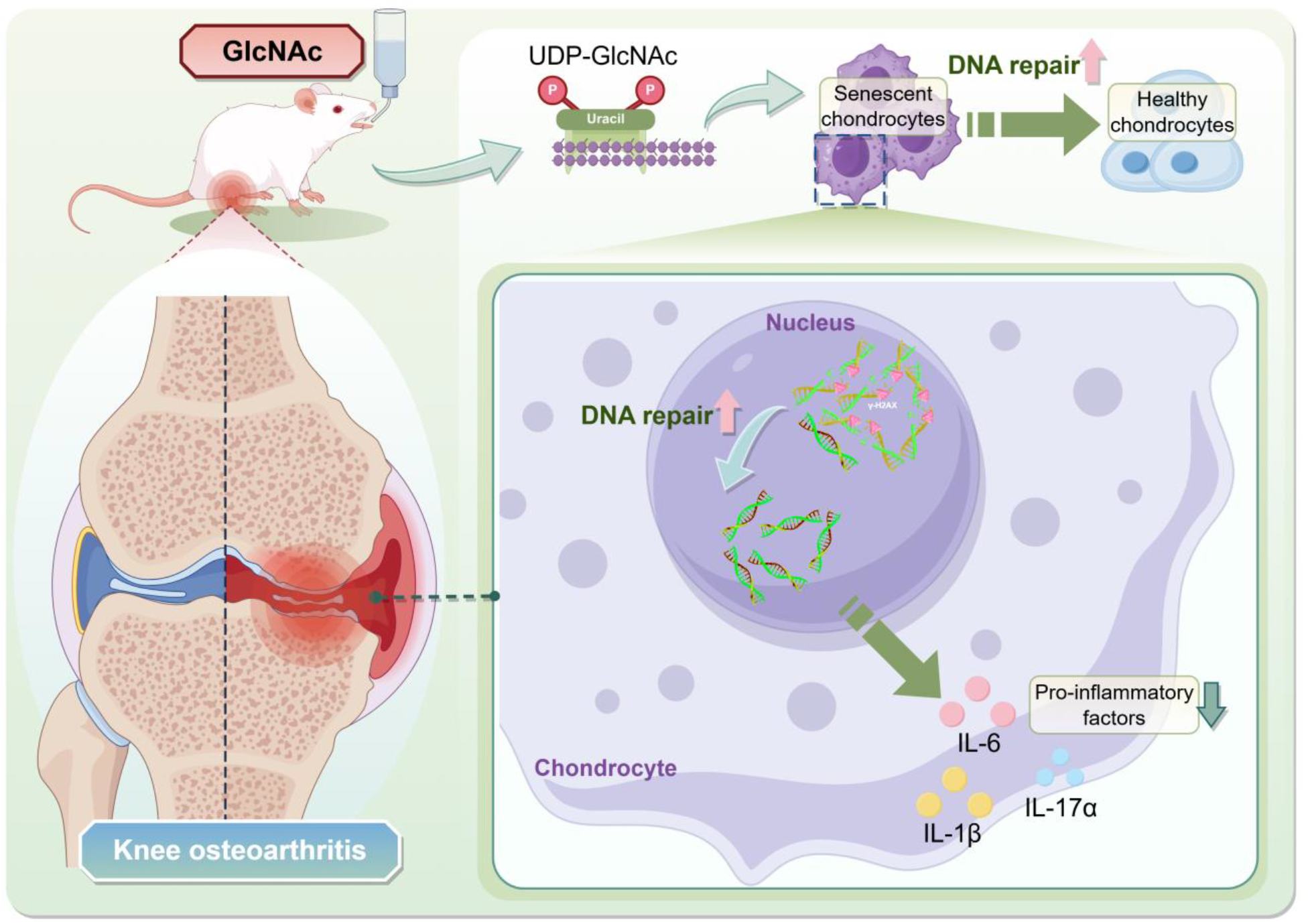
Schematic presents the role of GlcNAc in ameliorating knee osteoarthritis.

This study centrally contributes to elucidating the molecular mechanisms and signaling pathways underlying the differential therapeutic efficacy of GlcNAc and GlcN·HCl. The pharmacokinetic data demonstrated that GlcNAc achieved a 4.57-fold higher peak concentration of UDP-GlcNAc at 9 hours post-administration compared with GlcN·HCl, thereby establishing that the hexosamine biosynthetic pathway (HBP) flux critically determines downstream biological activity^[12]^. This finding directly addresses the research hypothesis by demonstrating that GlcNAc, as a direct precursor, bypasses the rate-limiting enzymatic step catalyzed by glucosamine-6-phosphate synthase (GFAT) and is more efficiently converted to the active metabolite UDP-GlcNAc, which serves as the obligatory substrate for glycosaminoglycan (GAG) synthesis in articular cartilage^[13]^. The transcriptomic analysis revealed that GlcNAc upregulated 2,808 DEGs compared with only 391 DEGs induced by GlcN·HCl, and showed significant enrichment of DNA repair and adaptive immune cell differentiation pathways. Using Random Forests machine learning, we identified four candidate DNA repair genes, *Hist1h2an, Poli, Ube2t*, and *Hells*, coupled with GSEA confirmation of homologous recombination repair pathway enrichment, this suggests that GlcNAc may protect chondrocytes from DNA damage-induced senescence or apoptosis, potentially through enhanced DNA repair in knee cartilage. These findings collectively establish that the superior therapeutic efficacy of GlcNAc results from its more efficient metabolic conversion to UDP-GlcNAc, which in turn activates both cartilage matrix synthesis and DNA repair pathways, thereby offering a twofold strategy for decelerating KOA progression that is fundamentally distinct from the symptomatic relief provided by conventional GlcN·HCl supplementation.

The functional and cellular behavioral implications of the genes in these findings substantially advance our understanding of chondrocyte biology under osteoarthritic stress and the regenerative capacity of articular cartilage. The downregulation of *Mmp3* and Adamts4 by GlcNAc treatment indicates a direct inhibitory effect on catabolic pathways in chondrocytes, as these enzymes are the principal proteases responsible for degradation of the cartilage extracellular matrix (ECM), particularly aggrecan and type II collagen^[14]^. The concomitant increase in cartilage area and reduction in scores across multiple independent histological grading systems, including Kellgren-Lawrence, ICRS, OARSI, Mankin’s, and synovitis scores, provides robust cross-validated evidence that GlcNAc shifts the balance from catabolism to anabolism in the joint microenvironment^[15]^. The upregulation of DNA repair genes suggests that GlcNAc may enhance the regenerative capacity of chondrocytes by maintaining genomic integrity under oxidative stress characteristic of the osteoarthritic milieu, thereby preventing the accumulation of senescent chondrocytes that contribute to disease progression^[16]^. This cellular protection mechanism is consistent with emerging evidence that DNA repair pathways are critical determinants of tissue homeostasis and that their impairment drives degenerative pathologies^[17]^. Furthermore, the observed reduction in pain-related neurochemical markers, *Pgp9*.*5* and *Ngf*, suggests that GlcNAc may modulate chondrocyte-derived neurotrophic signaling, thereby reducing nerve ingrowth and nociceptor sensitization within the damaged cartilage^[14]^. The functional changes in gene expression observed in this study therefore reveal that GlcNAc not only attenuates catabolic and inflammatory processes but also actively promotes cartilage repair through a coordinated upregulation of DNA repair genes and anabolic pathways, fundamentally altering our understanding of how metabolic precursors can be leveraged to modify the course of KOA rather than merely palliating its symptoms^[18]^. The identification of *Hist1h2an, Poli, Ube2t*, and *Hells* as GlcNAc-responsive DNA repair genes provides potential biomarkers for evaluating therapeutic efficacy and may represent novel molecular targets for disease-modifying osteoarthritis drug development, particularly given the established role of homologous recombination in maintaining genomic stability.

The immunological mechanisms elucidated in this study reveal a complex and multifaceted immunomodulatory role for GlcNAc in the KOA joint microenvironment that extends beyond simple immunosuppression. The immune infiltration analysis demonstrated that GlcNAc treatment led to a reduced abundance of pro-inflammatory M1 macrophages, regulatory T cells (Tregs), and CD4+ T cells, with concomitantly increased levels of monocytes, plasma cells, and CD8+ T cells. This pattern suggests that GlcNAc does not exert global immunosuppression but rather reshapes the immune landscape away from the chronic, low-grade inflammation characteristic of KOA pathogenesis, toward a more balanced and potentially reparative state. The observed reduction in M1 macrophages is particularly significant, as these cells are key producers of pro-inflammatory cytokines and matrix-degrading enzymes in the osteoarthritic joint^[19-21]^. The consistent reduction of IL-1β, IL-6, and IL-17α across ELISA, RNA sequencing, and FCM platforms provides corroborative, multi-platform validation of the anti-inflammatory effects of GlcNAc. Notably, IL-17α is particularly significant for its established role as a key cytokine in osteoarthritis pathogenesis and pain sensitization^[22]^. The observed elevation in CD8+ T cells, confirmed by FCM of bone marrow, may indicate a shift toward adaptive immune surveillance, potentially contributing to the clearance of senescent or damaged chondrocytes and thereby facilitating tissue repair^[23]^. This immunomodulatory profile is consistent with the broader concept that effective disease-modifying therapies for KOA must address the inflammatory microenvironment rather than solely targeting cartilage degradation, and it aligns with emerging evidence that GlcNAc-based compounds possess significant immunomodulatory and anti-inflammatory activities^[4, 9]^. Collectively, these immunological findings establish that GlcNAc exerts its therapeutic effects through a comprehensive remodeling of both innate and adaptive immune cell populations and their associated cytokine networks, providing a mechanistic foundation for its superior efficacy over GlcN·HCl in decelerating KOA progression.

We acknowledge the following limitations of this study: Despite the comprehensive multi-modal approach employed in this study, several methodological constraints merit acknowledgment. First, the relatively small sample size (n=8 per group) may compromise statistical power, particularly when detecting subtle transcriptomic or immunological differences. Next, while bulk RNA sequencing provides valuable pathway-level insights, it obscures cell-type-specific responses within heterogeneous cartilage tissue; single-cell resolution could more precisely delineate chondrocyte versus immune cell contributions. Then, the 60-day treatment window (127 days post-surgery) reflects only subchronic effects, leaving long-term efficacy, optimal dosing, and safety profiles unexamined. Finally, all findings are derived from a rodent model without human validation, and the translational relevance to KOA patients, including potential species-specific metabolic differences, remains to be established. Future studies addressing these gaps will be critical to advancement toward clinical application.

## Conclusion

This study demonstrates that GlcNAc demonstrates superior therapeutic efficacy compared with GlcN·HCl in a rat DMM model of knee osteoarthritis, mediated through multiple converging mechanisms. GlcNAc achieved a 4.57-fold higher conversion to the bioactive metabolite UDP-GlcNAc, and attenuated structural degeneration of knee cartilage by rebalancing the immune landscape away from a chronic inflammatory state and activating homologous recombination DNA repair pathways. Collectively, these findings position GlcNAc as a promising candidate that addresses both structural and symptomatic dimensions of KOA, warranting further clinical investigation.

## Ethics approval and consent to participate

This study was approved by the Institutional Animal Care and Use Committee (IACUC) of Sun Yat-Sen University (SYSU-IACUC-2026-B1855) and the Ethics Committee of Shenzhen Topbiotech Co. Ltd. (TOP-1PZ-GM250917). All experimental methods and procedures were conducted in accordance with the Animals in Research: Reporting of In Vivo Experiments Guidelines (https://www.nc3rs.org.uk/arrive-guidelines) and adhered to all relevant ethical regulations for animal testing and research.

## Acknowledgments

The authors thank OE Biotech Co. Ltd. (Shanghai, China) for bulk RNA sequencing, Topbiotech Co. Ltd. (Shenzhen, China) for experiments involving live rats, and the Instrumental Analysis Center of Shenzhen University for the assistance with micro-CT analysis. The schematic figures were manually drawn on Figdraw 2.0 of Home for Researchers.

## Author contributions

Conceptualization: Zilong Wang, Chunxia Zhang, ianzhi Zhang.

Methodology: Changze Song, Meilu Li, Xiangfei Qi, Huisheng Yuan.

Investigation: Changze Song, Meilu Li, Haohan Zhang, Junhao Chu, Xiaohan Wang, Yong Yang.

Visualization: Xiangfei Qi, Huisheng Yuan, Jinghan Zhang, Hongyin Huang, Zhenghao Li.

Funding acquisition: Zilong Wang, Changze Song.

Project administration: Zilong Wang, Chunxia Zhang, Tianzhi Zhang.

Supervision: Zilong Wang, Chunxia Zhang, Changze Song, Xiangfei Qi.

Writing – original draft: Changze Song, Meilu Li, Xiangfei Qi, Huisheng Yuan.

Writing – review & editing: Zilong Wang, Chunxia Zhang, Tianzhi Zhang.

## Data availability

All data are available in the main text or the supplementary materials.

